# Specific norovirus interaction with Lewis x and Lewis a on human intestinal inflammatory mucosa during refractory inflammatory bowel disease

**DOI:** 10.1101/2020.11.24.397125

**Authors:** Georges Tarris, Alexis de Rougemont, Marie Estienney, Maeva Charkaoui, Thomas Mouillot, Bernard Bonnotte, Christophe Michiels, Laurent Martin, Gael Belliot

**Author notes:** **Correspondence:** Dr. Gaël Belliot, Centre National de Référence des virus des gastro-entérites, Laboratoire de virologie, PBHU, CHU Dijon Bourgogne, 2 rue Angélique Ducoudray, BP37013, 21070 DIJON cedex, France. **Competing interest:** The authors declare no commercial affiliations or patent-licensing arrangements in the study design, the collection, analysis and interpretation of data, the writing of the report or the decision to submit the paper for publication that could be regarded as posing a conflict of interest concerning the submitted manuscript. **Specific author contributions:** Conceptualization: GT, LM, GB; Methodology: GT, GB; Investigation: GT, AdR, ME, GB; Resources: AdR, ME, MC, TM, CM, LM, GB; Data curation: MC, TM, CM; Validation: BB, CM, LM, GB; Writing original draft: GT, GB; Writing review & editing: AdR, BB; Funding acquisition: AdR, LM, GB; Supervision: LM, GB.

## Abstract

Inflammatory bowel disease (IBD), which includes Crohn's disease (CD) and ulcerative colitis (UC), is related to immunological and microbial factors with the possible implication of enteric viruses. We characterize the interaction between human noroviruses (HuNoVs) and blood group antigens in refractory CD and UC using HuNoV Virus Like Particles (VLPs) and histological tissues. Immunohistochemistry was conducted on inflammatory tissue samples from the small intestine, colon and rectum in 15 CD and 9 UC patients. Analysis of the regenerative mucosa of the colon and rectum revealed strong expression of sialylated Lewis a (sLe^a^) and Lewis x (sLe^x^) antigens, and HuNoV VLP binding in the absence of ABO antigen expression in both UC and CD. Competition experiments using sialidase, lectins and monoclonal antibodies demonstrated that HuNoV attachment mostly involved Le^a^ and to a lesser extent, Le^x^ moieties on regenerative mucosa in both UC and CD. Further studies will be required to understand the implications of specific HuNoV binding to regenerative mucosa in refractory IBD.

**IMPORTANCE:** Inflammatory Bowel Disease (IBD), including Crohn's disease (CD) and ulcerative colitis (UC), are progressive diseases affecting millions of people each year. Flare-ups during IBD result in severe mucosal alterations of the small intestine (CD), colon and rectum (CD and UC). Immunohistochemical analysis of CD and UC samples showed strong expression of known tumoral markers, Sialyl-Lewis a (CA19.9) and Sialyl-Lewis x (CD15s) antigens on colonic and rectal regenerative mucosa, concurrent with strong human norovirus (HuNov) VLP GII.4 affinity. Sialidase treatment and competition experiments using HBGA-specific monoclonal antibodies and lectins clearly demonstrated the implication of the Lewis a moiety, and to a lesser extent Lewis x, in HuNov recognition in regenerative mucosa of CD and UC tissues. Further studies are required to explore the possible implications of enteric viruses in the impairment of epithelial repair and dysregulation of inflammatory pathways during severe IBD.

## INTRODUCTION

Inflammatory bowel disease (IBD), which includes Crohn's disease (CD) and ulcerative colitis (UC), is a progressive gastrointestinal condition that affects millions of people worldwide. In Europe, IBD incurs healthcare costs totaling more than €5 billion every year (1). CD is characterized by mucosal ulcerations involving the entire intestinal tract, while UC only affects the colon and rectum. The two diseases have similar symptoms, including abdominal pain, fatigue, diarrhea and significant weight loss in some cases (2). Besides the fact that IBD is debilitating, patients have a greater likelihood of developing precancerous mucosal lesions (dysplasia) and cancer throughout the course of the disease. The etiology of IBD is currently not well understood, but the literature mainly suggests that it is due to autoimmunity combined with environmental factors and dysbiosis of the intestinal flora (3). IBD is also characterized by a reduction in glycosylation at the surface of the epithelium (4). It has been suggested that simplified glycan structures on the lumen of the intestinal cells might increase the odds of adventitious contact between enteric pathogens and the intestinal epithelium, therefore increasing the risk of inflammation (5). Infectious diseases are a recurring problem for people with IBD because treatment increases the risk of opportunistic infections (6). Among bacterial and viral enteric pathogens, human noroviruses (HuNoVs) are one of the most common causes of gastroenteritis in all age groups, and they are responsible for 18% of gastroenteritis cases worldwide. The genus *Norovirus* belongs to the *Caliciviridae* and is divided into seven genogroups (GI to GVII). GI and GII include the HuNoVs and are subdivided into eight (GI.1 to GI.8) and 22 genotypes (GII.1 to GII.22), respectively (7). Epidemiological surveys show that GII.4 HuNoV has been largely predominant in the world over the last two decades. There appears to be a strong correlation between the secretor phenotype and an increased susceptibility to norovirus infections in healthy individuals. So far, it is unclear whether HuNoV infections constitute a causal or an aggravating factor for IBDs (8), and there are conflicting reports about the increased detection rate of HuNoV in IBD patients. While the results of two epidemiological studies showed no correlation between HuNoV infection and CD (9), it has been hypothesized that an asymptomatic HuNoV infection may encourage a disruption in the intestinal flora, favoring the emergence of CD (10).

The natural ligands of the HuNoV are histo-blood group antigens (HBGAs). HBGAs are complex carbohydrate structures within ramified glycans that are part of mucin glycoproteins or glycolipids (11). An active *FUT2* gene encoding the type 2-fucosyltransferase is responsible for the expression of the A, B and H antigens in the small intestine and in the proximal colon, defining the secretor phenotype (12, 13). This phenotype contributes to the expression of Lewis b (Le^b^) and y (Le^y^) antigens, provided that the *FUT3* gene is active. For 20% of the European population, the *FUT2* gene is inactivated by a recessive nonsense mutation (14). The null allele defines the non-secretor phenotype, which is characterized by the absence of A, B, H, Le^b^ and Le^y^ antigens in mucosa and secretions. However, an active *FUT3* gene is responsible for the synthesis of Lewis a (Le^a^) and Lewis x (Le^x^) antigens in non-secretor individuals.

HBGAs are displayed at the surface of enterocytes in the small intestine, mainly in the duodenal mucosa. For all but the Le^a^ antigen, which has a pan-colonic distribution in both secretor and non-secretor individuals, HBGAs are found at the surface of the proximal colon but not the healthy distal colon or rectum (15-20).

The literature relative to the relationship between IBD and HBGA expression remains scarce. It seems that there is no obvious correlation between the distribution of ABO antigens and CD (21). The analysis of inflammatory colonic tissues for the presence of Sialyl-Lewis a (sLe^a^ also known as CA19.9) antigens during moderate and severe UC has shown that sLe^a^ molecules are absent during inactive and mildly active phases. However, the antigen is found in regenerative colonic epithelium in active UC, expressing similar markers to those observed in dysplasia (22). However, unlike dysplasia, sLe^a^ expression was found to be reversible in quiescent UC (19, 23). The increased risk of CD-related ileitis in individuals with non-secretor status was recently shown in a genetic linkage study of CD cases in the European population; this association has also been suggested for IBD (21, 24, 25). For UC, several polymorphisms of the *FUT3* gene could be involved in the genesis of the disease (26).

Microbial factors are also at play, and data show that IBDs could be related to changes in the bacterial composition of the microbiota, as exemplified by a decreased number of *Faecalibacterium prausnitzii* during flare-ups (27). The combination of these changes and the appearance of new pathogens could disrupt or exacerbate host immune response during colitis (28). However, little is known about the role of viruses. To date, we know that the bacteriophage composition changes during IBD – *Caudovirales* and *Microviridae* become predominant whilst there is a bacterial impoverishment (29). Still, we are unsure whether the observed increase in bacteriophages is an aggravating factor or a consequence.

Data from murine models show that IBD-triggering factors include a combination of immunological and genetic elements as well as the occurrence of infection. The use of cultivatable murine norovirus (MNV) in mice has shown that mucosal inflammation of the intestine is induced by viral infection if the mice are unable to produce the anti-inflammatory IL-10 cytokine (30). Additionally, as a proof of principle, Cadwell *et al*. have shown that the combination of genetic background and viral infection could induce structural changes in certain intestinal cells (i.e. Paneth cells), excessive inflammatory response, and Crohn's disease-like ileal disorder (31). The data from murine models suggest that similar triggering and/or aggravating factors may emerge during viral infection in humans as well. That being said, the role of the widely spread HuNoVs is still largely unknown in the context of IBD, much like the role of the expression of the HBGA natural HuNoV ligands. In the present study, we determined the putative interaction between HuNoVs and intestinal mucosa during IBD flare-ups. We focused on characterizing HBGA expression and HuNoV interaction at the surface of inflammatory tissue in samples from individuals diagnosed with UC or CD.

## RESULTS

### Histological observations and FUT2 genotype

We analyzed 19 samples of resected tissue from CD patients, including samples from the ileum (N=7), proximal colon (N=3), transverse colon (N=4) and sigmoid (N=2). We also analyzed 9 samples of resected tissue from UC patients, including samples from the sigmoid colon (N=6) and rectum (N=3) (Table 1).

**Table 1:** Cohort used for the study

| Sample<br>a, b | Age<br>(gender) <sup>c</sup> | Pathology <sup>d</sup> | Anatomical site | HES stain <sup>e</sup> | Blood<br>group | FUT2<br>genotype <sup>f</sup> |
| --- | --- | --- | --- | --- | --- | --- |
| 1 | 55 (M) | CD | Ileum | QM (80%) / RM (20%) | O | Sec/Sec |
| 2 | 25 (F) | CD | Ileum | QM (100%) | A | Sec/Sec |
| 3 | 34 (F) | CD | Ileum | QM (80%) / RM (20%) | AB | Sec/Sec |
| 4 | 22 (F) | CD | Ileum | QM (100%) | A | Sec/Sec |
| 5 | 25 (F) | CD | Ileum | QM (100%) | A | Sec/Sec |
| 6 | 29 (M) | CD | Ileum | QM (30%) / RM (70%) | O | Sec/Sec |
| 7 | 49 (F) | CD | Ileum | QM (90%) / RM (10%) | A | Sec/Sec |
| 8 | 46 (M) | CD | Proximal Colon | QM (100%) | O | Sec/sec |
| 9 | 42 (F) | CD | Proximal Colon | QM (100%) | A | Sec/Sec |
| 10 | 36 (F) | CD | Proximal Colon | QM (100%) | A | Sec/sec |
| 11 | 32 (M) | CD | Transverse Colon | QM (80%) / RM (20%) | A | Sec/sec |
| 12 | 24 (F) | CD | Transverse Colon | QM (100%) | B | Sec/Sec |
| **13** | 26 (F) | CD | Transverse Colon | QM (50%) / RM (50%) | A | Sec/sec |
| 14 | 30 (F) | CD | Transverse Colon | QM (100%) | A | Sec/sec |
| 15 | 22 (M) | CD | Sigmoid | QM (100%) | O | Sec/sec |
| 16 | 50 (F) | CD | Sigmoid | QM (90%) / RM (10%) | A | Sec/Sec |
| 17 | 33 (M) | UC | Sigmoid | QM (100%) | A | Sec/Sec |
| 18 | 42 (M) | UC | Sigmoid | QM (90%) / RM (10%) | O | Sec/sec |
| 19 | 30 (F) | UC | Sigmoid | QM (100%) | O | Sec/Sec |
| 20 | 26 (F) | UC | Sigmoid | RM (100%) | B | Sec/sec |
| 21 | 47 (M) | UC | Sigmoid | QM (30%) / RM (70%) | O | Sec/sec |
| **22** | 40 (F) | UC | Sigmoid | RM (100%) | A | Sec/sec |
| 23 | 32 (F) | UC | Rectum | QM (100%) | A | Sec/sec |
| 24 | 42 (F) | UC | Rectum | RM (100%) | O | Sec/sec |
| 25 | 58 (M) | UC | Rectum | QM (60%) / RM (40%) | AB | Sec/sec |

<sup>a</sup> samples 13 and 22 whom the samples were used for the competition experiments are indicated in bold.
<sup>b</sup> samples 7 and 16 correspond to two different surgeries for the same patient at 8-month interval
<sup>c</sup> age of the patient at the time of the surgical procedure is given in years. M : male. F : female.
<sup>d</sup> CD : Crohn's Disease. UC : Ulcerative Colitis
<sup>e</sup> Percentage of mucosal surface area for each patient sample. QM : Quiescent Mucosa. RM : Regenerative Mucosa.
<sup>f</sup> Sec/sec : heterozygous secretor. Sec/Sec : homozygous secretor.

Preliminary histological observations ruled out discernable features of dysplasia. The mucosa samples were classified as quiescent mucosa (QM) or regenerative mucosa (RM) (Table 1). QM was characterized as mildly inflamed or healed mucosa without major architectural alterations. RM was characterized as severely inflamed mucosa with marked architectural alterations and regenerative epithelium. We determined the percentage of QM and RM for each tissue sample. In 9 CD and 3 UC samples, only QM was observed (i.e. QM 100%). In 7 CD and 3 UC samples, both QM and RM were observed. Finally, in 3 UC samples (patients 20, 22 and 24) the tissue was exclusively regenerative (i.e. RM 100%). Additionally, the nuclear proliferative marker Ki-67 antigen was detected in proliferative niches located in the basal crypts compartment, with low levels detected in the tissue defined as regenerative mucosa (32, 33) (Fig. S1). FUT2 genotyping was performed in order to identify secretor and non-secretor individuals. The entire cohort belonged to the secretor genotype, including 2 homozygous and 7 heterozygous UC patients, and 9 homozygous and 6 heterozygous CD patients. No statistically significant genotype distribution was observed within the cohort (Table 1).

### Norovirus binding specificity

Baculovirus-expressed synthetic virus-like particles (VLPs) were used to mimic HuNoV capsid behavior. Binding assays on healthy duodenal sections displayed VLP attachment on the apical surface of enterocytes, as previously described in physiological conditions (34) (Fig. 1). VLP binding was abolished after sodium periodate treatment, demonstrating the involvement of carbohydrates. The absence of binding using ΔD373 VLPs confirmed that the attachment of GII.4 VLPs was HBGA-specific (Fig. 1). This observation was correlated with a strong reduction in VLP binding capacity following α1,2 fucosidase treatment. The next objective was to determine whether there were alterations in HBGA expression in CD and UC tissues, and thereby in norovirus VLP binding.

**Fig. 1:**
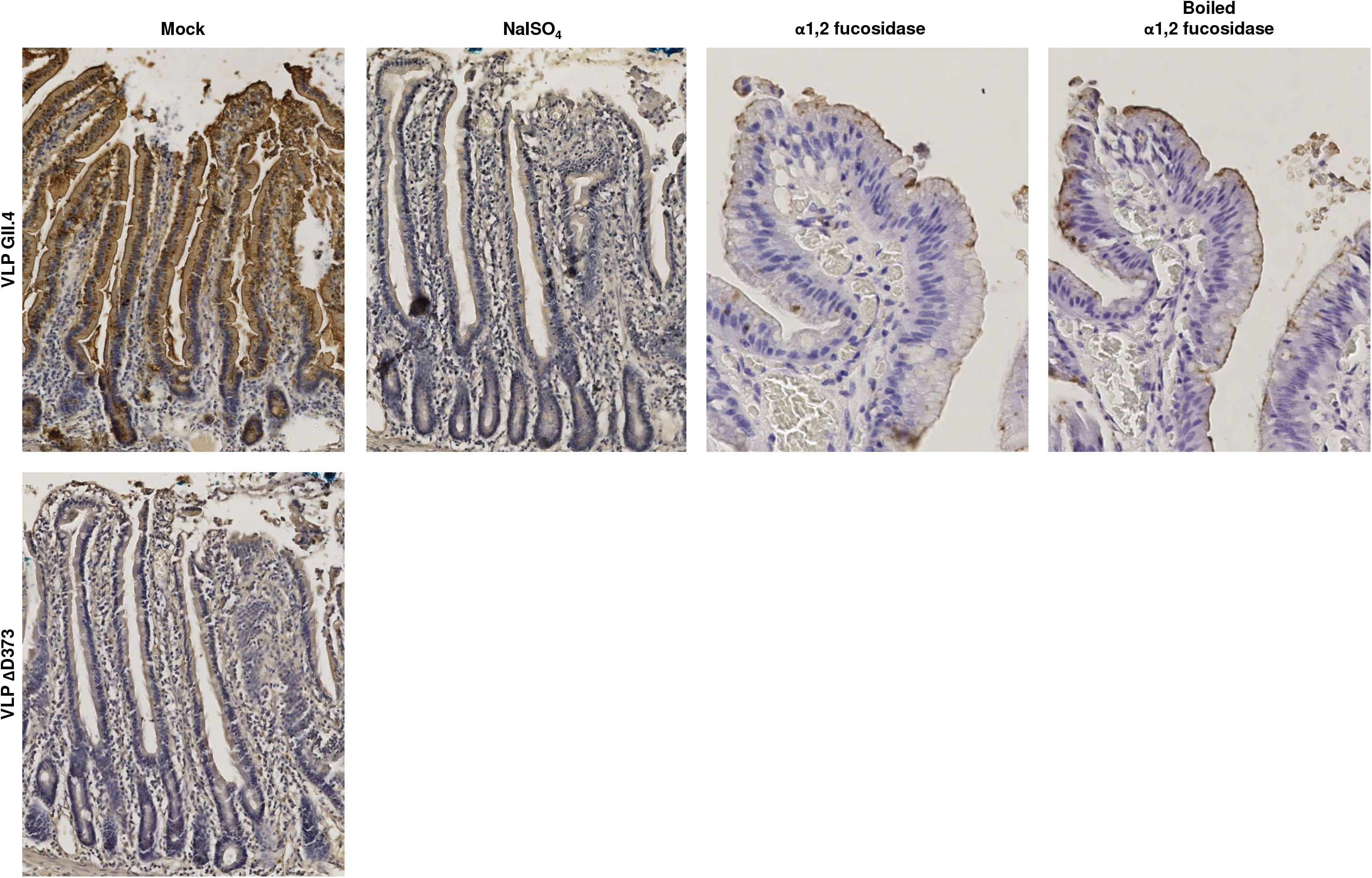
VLP binding specificity demonstrated in a healthy duodenal sample from a blood group A patient who underwent duodenopancreatectomy. Slides were pretreated with either 50mM sodium periodate (NaIO4), α1,2 fucosidase (boiled or cold) before incubation with HuNoV GII.4 VLP. Mutated ΔD373 GII.4 VLPs were used as negative controls. For this experiment and the following, VLP binding was detected with GII.4 VP1-specific MAb labeled with peroxydase. Peroxydase activity was detected by colorimetry using H2O2 and 3,3’-diaminobenzidine (DAB) giving a brown staining. Mock and pretreatments are indicated above each image.

### Crohn’s Disease

All the samples from CD patients contained QM, and 7 of these samples also included RM (10 to 70% of the total mucosal surface) (Supplemental Table 1). Ileal samples strongly expressed ABO antigens, which colocalized with VLP binding for RM and QM (Fig. 2 and 3). In RM, the distribution of Lewis antigens (i.e. Le^a^, sLe^a^, Le^x^ and sLe^x^) was pan-mucosal. In QM, the presence of Lewis antigens was somewhat contrasted: Le^a^, sLe^a^ and Le^x^ were only found in goblet cells, and their distribution varied considerably from one patient to another (10 to 80% of goblet cells). sLe^x^ expression was limited to 5 to 10% of the basal crypts in QM.

**Fig. 2:**
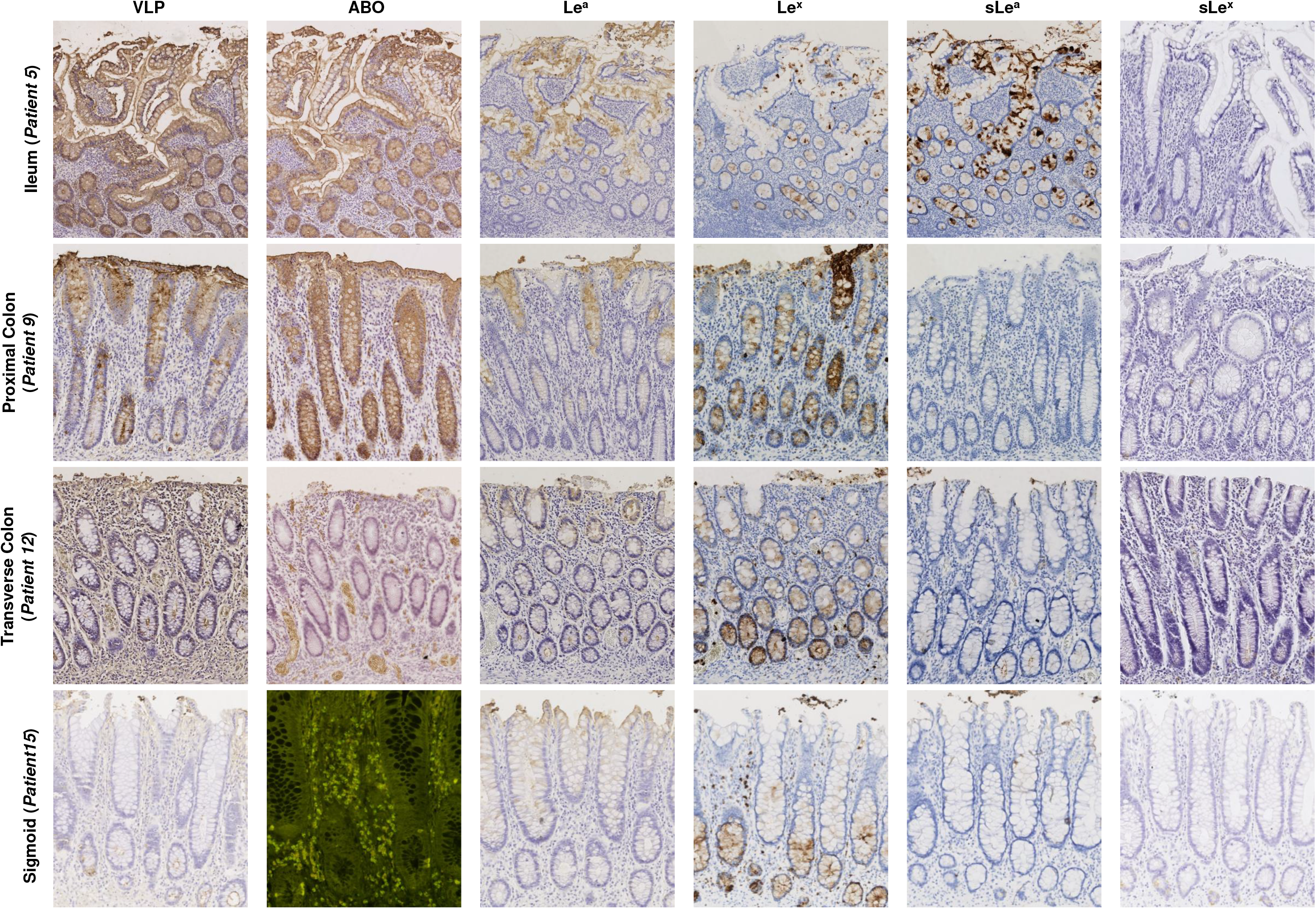
HBGA detection and HuNoV binding in CD ileum (patient 5), proximal colon (patient 9), transverse colon (patient 12) and distal colon (patient 15) showing quiescent mucosa. For this figure and the next two figures, blood group A, Le^a^, Le^x^, sLe^a^ and sLe^x^ antigens were detected with.specific Mabs in colorimetric assay using DAB. α1,2 fucose moiety characterizing H antigen was detected using FITC-conjugated UEA-I lectin. Detected antigens are indicated above each column. Patients (in parentheses) and tissue samples are indicated at the left of each row.

**Fig. 3:**
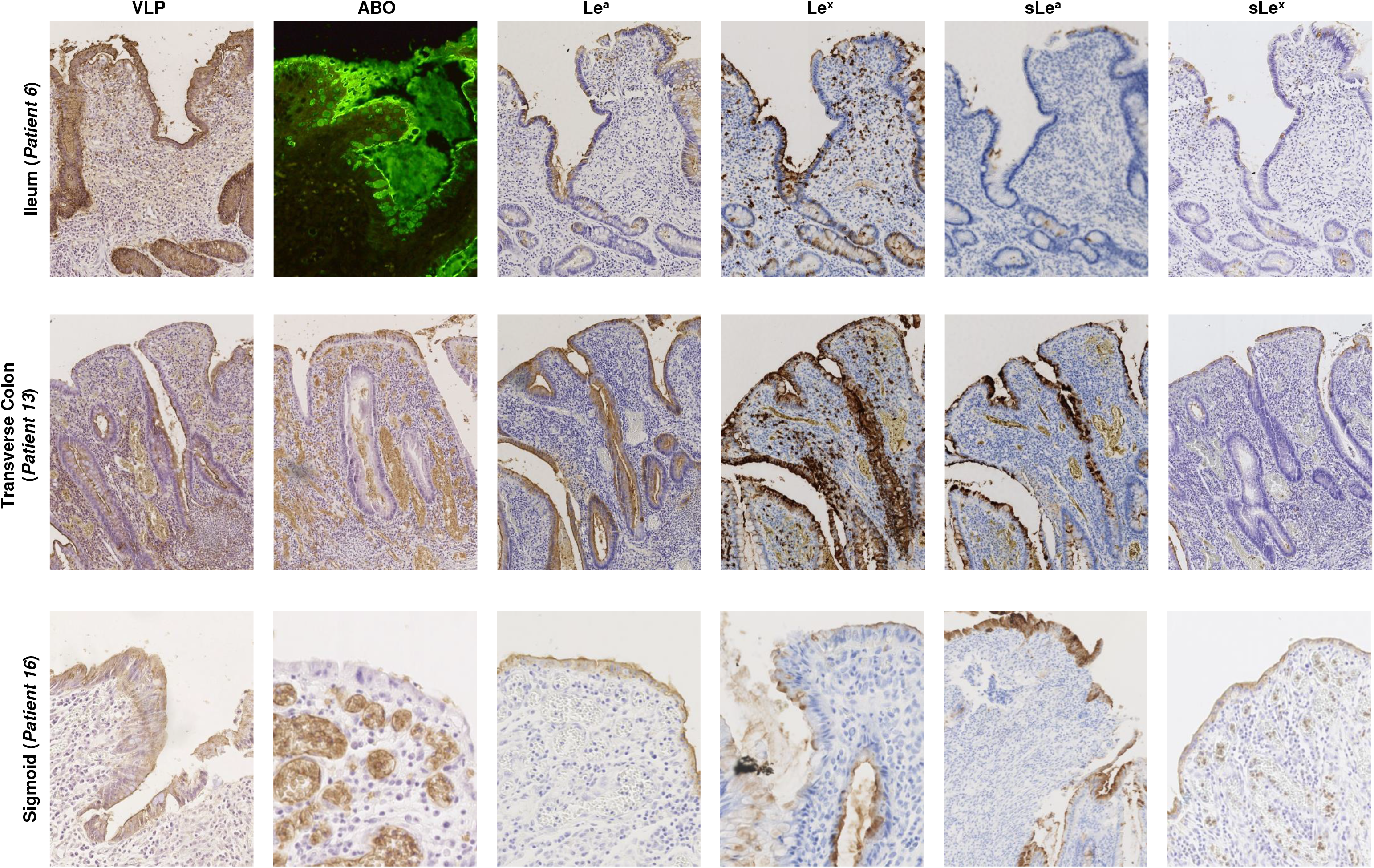
HBGA detection and HuNoV binding in CD ileum (patient 6), transverse colon (patient 13), and sigmoid (patient 16) showing regenerative mucosa. Detected antigens are indicated above each column. Patient numbers (in parenthesis) and tissue samples are indicated on the left of each row.

Proximal colon samples showed pan-mucosal distribution of the ABO antigens, which correlated with pan-mucosal attachment of HuNoV VLPs. Le^a^ and Le^x^ antigens were strongly expressed in goblet cells, while only 1 to 5% of goblet cells expressed sLe^a^ antigens. In ileal samples characterized as QM, the sLe^x^ antigen was expressed in only 1 to 5% of basal crypts (Fig. 2). In transverse colon samples, ABO antigens were weakly detected in RM (0 to 0.5%) and absent in QM (Fig. 3). Surprisingly, we did observe VLP binding to basal crypts and goblet cells in QM, while in RM there was a marked pan-mucosal distribution of VLP binding despite the absence of ABO antigens. We also observed a strong expression of Lewis and sialylated Lewis antigens in all RM mucosa. Le^a^ expression was restricted to goblet cells in QM (30 to 80%) except for patient 14, who showed pan-mucosal expression of the antigen. Antigen expression in goblet cells was also observed for Le^x^ (10 to 70%) and sLe^a^ (5 to 30% of goblet cells). SLe^x^ expression was restricted to 5% of the total mucosa, corresponding to the basal crypts.

For sigmoid samples, ABO antigens were absent in QM and poorly expressed in RM (mean 0.25%). VLP binding was observed in all RM (100%), contrasting with QM which displayed poor VLP attachment (mean 20%) (Fig. 3). We observed strong pan-mucosal expression of the Lewis and sialylated Lewis antigens in RM. In QM, Le^a^ and sLe^a^ were only expressed in goblet cells, accounting for 10 to 50% and 5 to 10% of the cells, respectively. Le^x^ and sLe^x^ expression accounted for 20 to 30% and 5% of the cells, respectively, and were mainly restricted to basal crypts.

### Ulcerative Colitis

The samples from the 8 UC patients contained a combination of QM and RM. Patient 17 had only QM and patient 22 had only RM. For sigmoid and rectal samples, strong VLP attachment was observed in RM while mucosal ABO expression remained absent or patchy (patients 18 and 25) (Fig. 4). VLP binding was concomitantly associated with strong expression of Le^a^ (50 to 100% of mucosa), sLe^a^ (50 to 100% of the mucosa), Le^x^ (80 to 100% of mucosa) and sLe^x^ (80 to 100% of mucosa) (Supplemental table 2). In sigmoid and rectal samples with QM, VLP binding remained localized to 20 to 30% of basal crypts and also up to 60% of goblet cells (patient 18). There was a large distribution of Le^a^ (50 to 80%) and sLe^a^ (30 to 70%) in goblet cells, while expression of Le^x^ and sLe^x^ was somewhat lower in the same samples. In patients 17, 18 and 19, Le^x^ and sLe^x^ antigens were found in 10 to 30% and 5 to 20% of basal crypts, respectively. In patient 21, Le^x^ and sLe^x^ antigen expression was restricted to 30 and 10% of goblet cells, respectively. For QM in the sigmoid, VLP binding was scattered in 5 to 10% of basal crypts (Fig. 4). Contrary to the tissues described above, there was almost no expression of Le^a^ and sLe^a^. Inversely, 20 to 40% of the basal crypts expressed Le^x^, while sLe^x^ expression was only 5 to 10%.

**Fig. 4:**
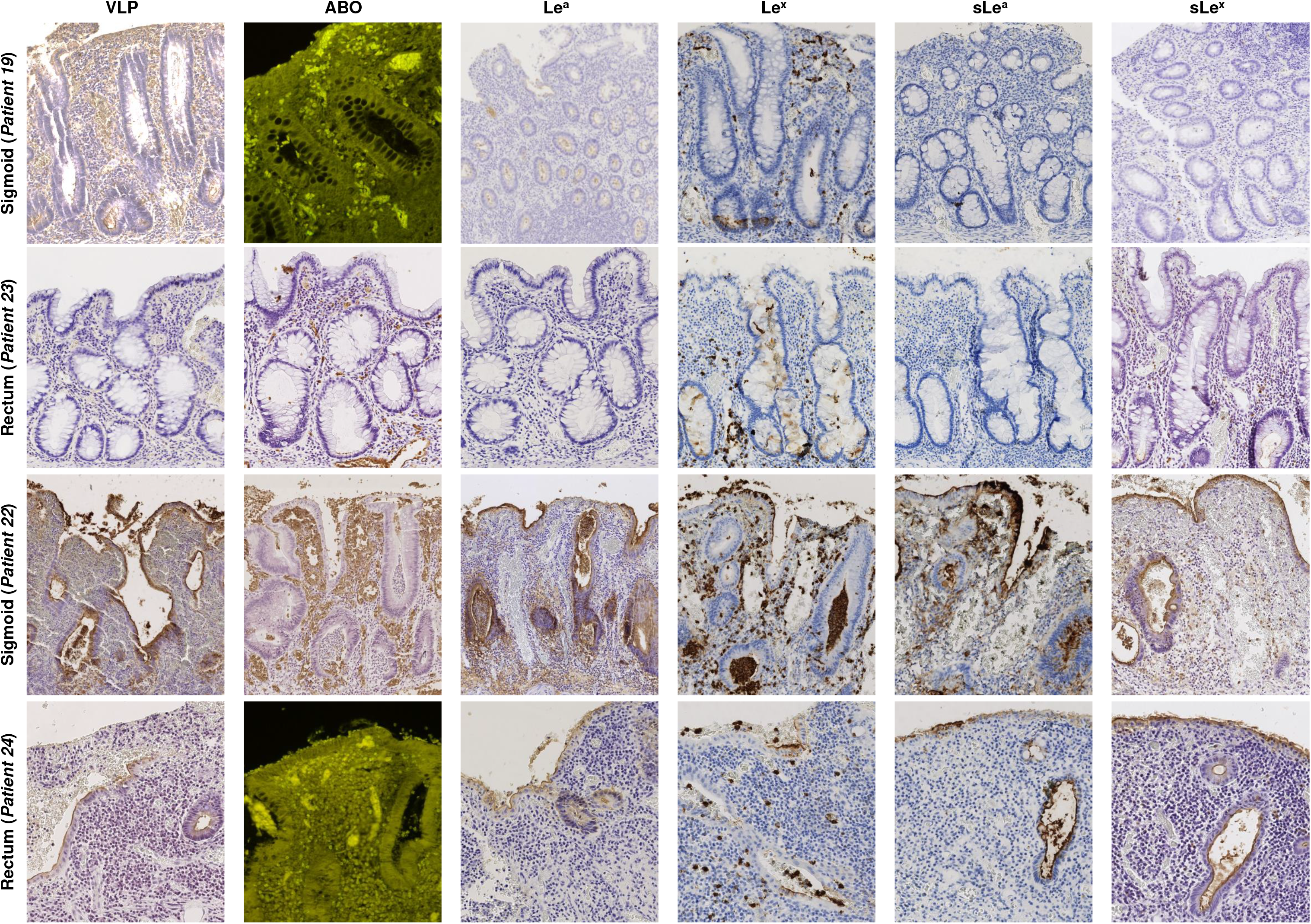
HBGA detection and HuNoV binding in UC colon (patient 19) and rectum (patient 23) with quiescent mucosa (QM), and sigmoid (patient 22) and rectum (patient 24) with severe inflammation and regenerative mucosa (RM). The detected antigen is indicated above the column panel. QM, RM, patients and tissue samples are indicated on the left of each row.

In summary, our analysis of refractory UC and CD samples shows discordant patterns of VLP binding during epithelial regeneration in the transverse colon, sigmoid and rectum, with poor ABO antigen expression. In addition, marked VLP attachment in RM correlated with a strong expression of Lewis antigens from the ileum to the rectum. The CD samples from the ileum and proximal colon strongly expressed ABO antigens and showed marked VLP attachment in QM. QM from the transverse colon to the rectum did not express ABO antigens, and VLP attachment was almost exclusively located in goblet cells and basal crypts. Lewis and sialylated Lewis antigens were also present in goblet cells and basal crypts in QM. Our data suggest that other ligands might be responsible for HuNoV binding, especially in RM where ABO antigens were absent. Previous in vitro studies showed that GII.4 HuNoV specifically recognizes the sLe^x^ antigen, with no in vitro recognition of Le^x^ or sLe^a^ (35). We thus hypothesized that sLe^x^ and/or other Lewis antigens are responsible for HuNoV binding by inflammatory tissues (36).

### Characterization of norovirus attachment on regenerative mucosa

#### Ulcerative colitis

One macrodissected sigmoid colon sample (patient 22; Table 1) was selected for further characterization of HuNoV attachment (Fig. 5). In a preliminary experiment, the removal of sialic acid moieties from the distal colon did not inhibit VLP attachment to inflammatory tissues, suggesting that the sialic acid moiety from sLe^x^ and sLe^a^ were not involved in HuNoV recognition (Fig. 5). Because ABO antigens are the main natural ligands for norovirus binding, competition experiments using specific lectins verified the role of the A and H antigens in VLP attachment. *Helix pomatia* and UEA-1 are specific lectins of the A and H antigens, respectively. Control assays performed on a duodenal biopsy sample from patient 22 showed that the preincubation of sections with both lectins together abolished HuNoV GII.4 VLP binding, indicating that both lectins efficiently blocked VLP attachment to the A and H antigens (Fig. S2). Similarly, both lectins were incubated with histological sections of the colon harboring inflammatory areas. Binding assays showed that VLPs did interact with regenerative epithelium despite inhibition of the A and H antigen binding activity, proving that other ligands were also involved in HuNoV attachment. Furthermore, no VLP binding to HBGAs was observed when using ΔD373 VLPs, suggesting that HBGA or HBGA-like antigens were still involved (data not shown) (37).

**Fig. 5:**
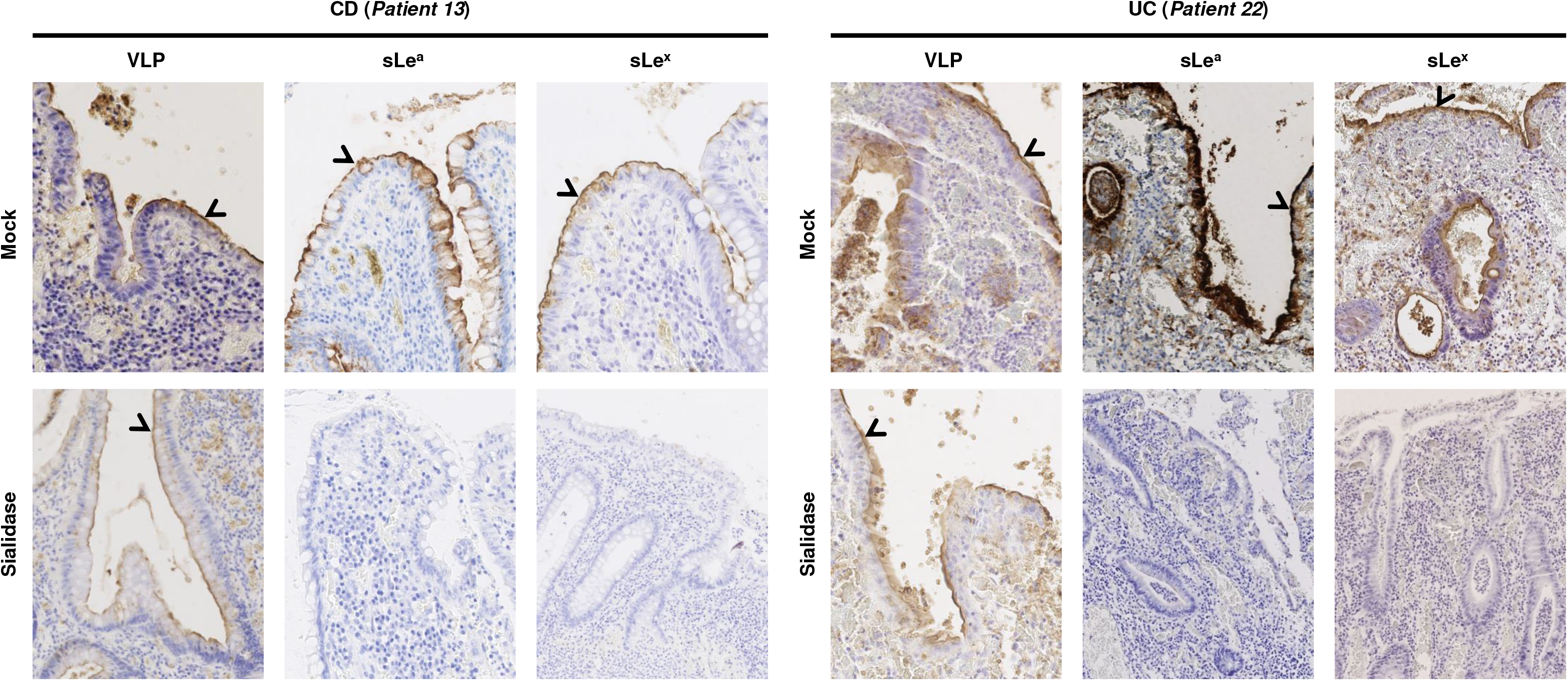
Role of the sialic acid moiety into HuNoV attachment. Attached VLP to mucosa and presence of sLe^a^ and sLe^x^ were detected in CD (patient 13) and UC (patient 22) macrodissected tissue samples following sialidase treatment. The efficacy of the sialidase treatment for the removal of the sialic acid was controlled by the absence of immunostaining following incubation with sLe^a^ and sLe^x^-specific Mabs. Mock and sialidase treatments are indicated on the left of each row. Detected antigens are indicated above each column panel. Specific staining is indicated with an arrowhead.

In order to determine the role of each antigen in HuNoV recognition, tissue sections were incubated with HPA and UEA-I lectins to efficiently suppress putative A and H antigen binding activity on regenerative mucosa. *Lotus tetragonolobus* (LTL) and *Aleuria aurantia* (AAL) lectins were also used to inhibit VLP attachment. LTL can specifically recognize Le^x^ antigens if the sialic acid moiety is absent (38). Therefore, tissue sections were pretreated with sialidase, then preincubated with HPA and UEA-I lectin, and finally incubated with LTL (Fig. S3). The marked VLP attachment on inflammatory areas suggested that the Le^x^ antigen did not play an important role in HuNoV recognition. AAL specifically recognizes α1-2, α1-3, α1-4 and α1-6 fucose moieties and can therefore recognize Le^a^ and Le^x^ antigens (39, 40). The AAL lectin completely inhibited VLP attachment, whereas the use of boiled lectin as a negative control was not able to inhibit HuNoV recognition (Fig. S4). Our data suggests that α1-3 (Le^a^) and/or α1-4 (Le^x^) fucose were the main ligands involved in HuNoV binding in regenerative mucosa in UC. To pursue further, the sections were incubated either with Le^x^-specific or Le^a^-specific antibodies, or both, after sialidase treatment and preincubation with HPA and UEA-I lectins (Fig. 6). Weaker VLP binding still occurred with Le^x^-specific antibodies, while VLP binding was totally abolished when Le^a^-specific antibodies were incubated alone or in addition to Le^x^-specific antibodies (Fig. 6). We found that the Le^a^ antigen and to a lesser extent the Le^x^ antigen (alone or as part of sLe^a^ and sLe^x^ molecules) were responsible for the specific recognition of HuNov GII.4 VLP on regenerative mucosa in UC.

**Fig. 6:**
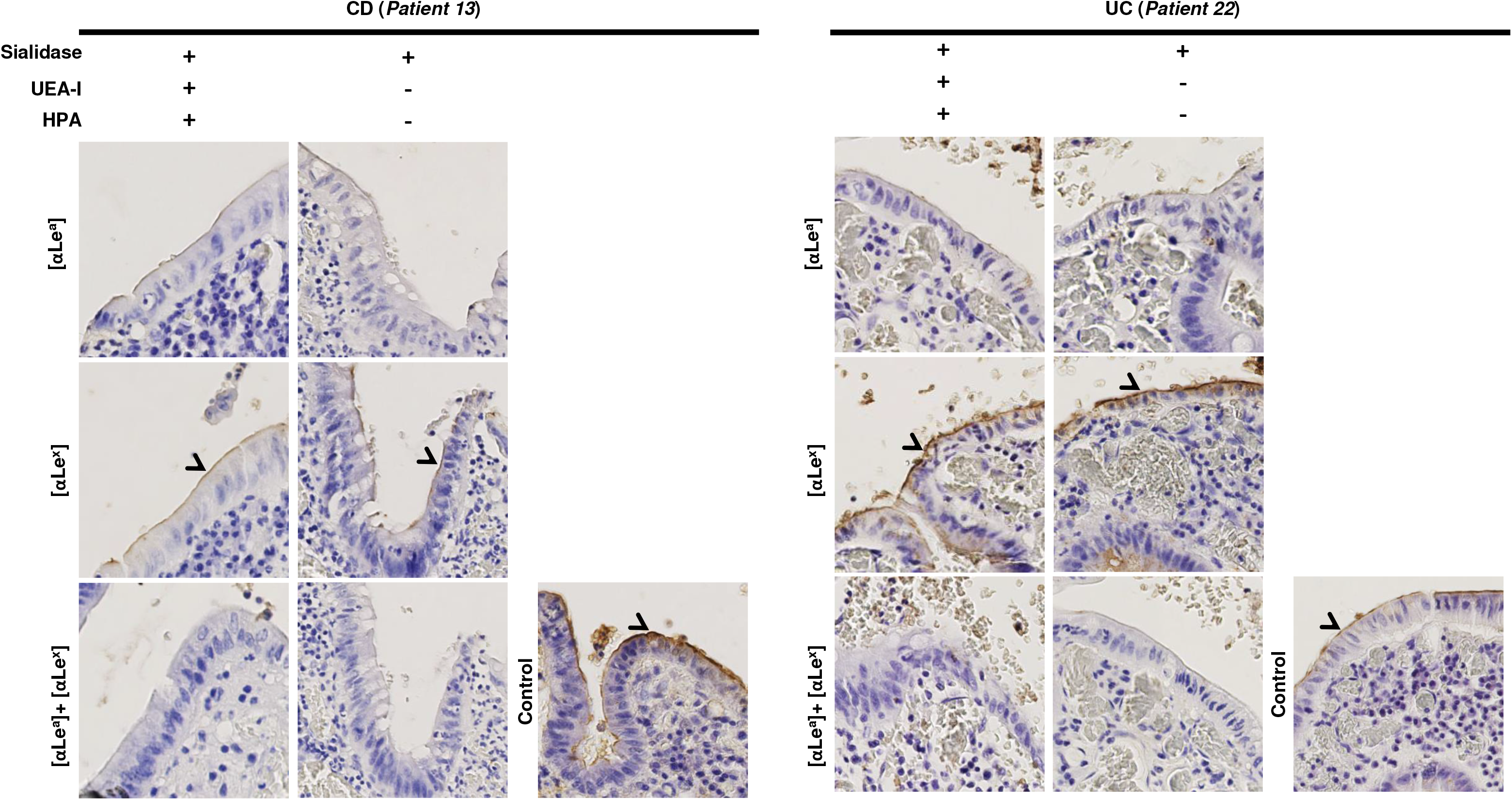
Role of the Lewis antigens into HuNoV specific attachment to pathological CD (patient 13) and UC (patient 22) mucosae. Tissues were first preincubated with a combination of sialidase and lectins (UEA-I and HPA). Presence or absence of treatment is indicated above the column panel by positive and minus signs, respectively. The objective was then to specifically inhibit VLP with Lewis-specific Mabs. Slides were first pretreated with specific mAbs against Le^a^ (αLe^a^), Le^x^ (αLe^x^), or both combined before HuNoV GII.4 VLP incubation. The Mabs used for the experiments are indicated in bracket at the left side of each row. For the control, VLPs were directly incubated on tissue without any pretreatment with sialidase, lectins or Mabs. VLP attachment is indicated by arrowheads and magnified areas correspond to dashed boxes.

#### Crohn’s disease

Data obtained from the CD group showed that regenerative mucosa strongly expressed sLe^a^ and sLe^x^ ligands, in addition to specific HuNoV GII.4 VLP binding without significative ABO antigen expression. Selective inhibition using specific antibodies and lectins helped to characterize HuNoV GII.4 VLP binding in refractory CD.

Competition experiments were performed using a macrodissected transverse colon sample for further characterization of VLP binding (patient 13, Table 1). Sialidase treatment failed to inhibit HuNov GII.4 VLP binding, again suggesting that the sialic acid moiety was not involved in HuNov GII.4 VLP recognition (Fig. 5). After sialidase pretreatment and incubation with HPA and UEA-I lectins, tissue sections were incubated with Le^a^-specific and Le^x^-specific antibodies (individually and in combination), which abolished HuNoV GII.4 VLP binding (Fig. 6).

Our results show that HuNoV GII.4 VLPs specifically recognize Le^a^ and sLe^a^ and to a lesser extent Le^x^ antigens in regenerative mucosa during both refractory CD and UC. On the contrary, we observed poor HuNoV GII.4 VLP binding in quiescent and healthy colonic and rectal mucosa, without significant ABO expression.

## DISCUSSION

The immunological aspects of IBD and the dysbiosis of the intestinal flora have largely been documented in the literature, yet little is known about the interactions between viral enteric pathogens and the intestinal tract when it is affected by IBD. In both CD and UC, we observed that Le^a^ and Le^x^ antigens and their sialylated counterparts were specifically expressed by inflammatory and regenerative tissues from all patients, especially in RM. The presence of these ligands was striking, especially in the colon where they are absent in physiological conditions. Our data can be explained by the fact that regenerative and inflammatory tissues are characterized by a capacity to express a large panel of molecules; this capacity is lost once cell differentiation is achieved. Since HBGAs are natural ligands for HuNoVs, the objective of our study was to determine the interaction between HuNoVs and pathological mucosae from the intestine, colon and rectum of patients with IBD.

One recent study has clearly shown that the small intestine is the main replication site for HuNoVs as a result of highly expressed HBGAs (41), while it is assumed that HuNoVs do not replicate in the colonic epithelium of healthy adults (42). In our preliminary experiments, we did indeed observe that HuNoV VLPs specifically recognized HBGA molecules on duodenal cells, as described previously (34). On the contrary, VLP did not bind at the surface of epithelial cells in the healthy tissue of the colon where HBGAs are not normally expressed. Analysis of pathological tissues from the colon and rectum of CD and UC patients showed structurally disorganized cells from regenerative epithelium on the luminal side, and, unlike in healthy tissue, we observed a strong attachment of GII.4 VLPs. Additional experiments using GII.3 and GII.17 VLPs also showed that GII.17 did specifically bind to regenerative and inflamed areas while GII.3 did not, suggesting that HuNoV binding in the context of IBD was genotype-specific (Fig. S5). At the cellular level, VLP attachment occurred on the surface of the regenerative mucosa expressing sLe^a^ and sLe^x^ in the absence of histological features characterizing precancerous lesions. The binding of HuNoV to sLe^x^ has previously been characterized in vitro using GII.4 HuNoV VLP, but the biological significance of this process remains unclear (35). Rydell *et al* suggested that there is a “sialic pathway” for norovirus attachment in addition to the established “α1-2 fucose” pathway characterizing HuNoV-HBGA interaction. Our data showed that Le^a^ and Le^x^ antigens, alone or associated with sialic acid moiety, were responsible for HuNoV capsid recognition on inflammatory and healing tissues during UC and CD flare-ups. Future research should thus focus on the fate of HuNoVs after they bind to Le^a^ and Le^x^ ligands.

In healthy intestines, HuNoV replication is generally thought to take place mainly in the enterocytes where HBGAs are abundantly expressed (43). Although viral attachment is followed by the internalization of the particles in the cells before their replication, we do not know whether HuNoV attachment triggers their internalization and replication in regenerative cells. We are therefore left wondering about the physiological consequences of HuNoV replication, specifically in inflammatory mucosa from the colon in IBD. Since the etiology of IBD is complex and multifactorial, the role of enteric viruses remains unclear. This is especially true for the dysbiosis seen in IBD microbiota (29). We could hypothesize that this unusual attachment might be involved in the disruption of the intestinal flora. However, although we previously demonstrated that HuNoV VLPs could attach to injured mucosa from the colon and rectum, cells from healing and inflammatory tissues might obstruct HuNoV replication, even after successful attachment. That being said, the newly formed virus-HBGA complex might activate the immune system and exacerbate the inflammatory process. In the future, it will be important to determine whether the presence of viral enteric pathogens and their interaction with enteric cells is correlated with the inflammatory response observed in IBD. This would require the histological analysis of IBD patients suffering from infection with HuNoV gastroenteritis or other enteric viruses such as group A rotaviruses, which also recognize HBGA ligands (44, 45).

One shortcoming of our study is the limited number of UC and CD samples, limiting the robustness of our statistical analysis. In addition, the experiment showing that Le^a^ and Le^x^ antigens were involved in the recognition of HuNoV was only conducted on the samples from two patients. Only severe cases of IBD were included in the study in compliance with the recommendation of the French national ethical committee, which explains only 27 patients were included in our study. Additional studies are therefore needed to pursue our research, provided there is enough available biological material. That being said, it is worth mentioning that inflammatory areas were characterized for all UC and CD patients of the study and showed the presence of Le^a^ and Le^x^ antigens. This observation strongly suggests that both antigens might be involved into NoV recognition on inflammatory areas. The other limitation of the study is the use of VLPs that might not entirely reflect biological properties of native HuNoV particles. Even though VLPs are not infectious like native particles, they have been essential for demonstrating the role of HBGAs as HuNoV natural ligands (34). Therefore, the specific binding of VLPs or native HuNoV on inflammatory mucosa is not necessarily followed by viral replication However, virus attachment may trigger activation of the immune system.

Further study of the microbiota in combination with epidemiological data will help to unravel the precise role of common viral enteric pathogens during IBD. Recent findings regarding HuNoV replication in human intestinal enteroids (HIEs) and organoids (HIOs) are promising (46, 47). The use of organoids derived from IBD patients might be useful to study specific virus-host interactions and genetic responses, as described previously (48).

From a clinical point of view, epidemiological studies have shown that opportunistic HuNoV infections require special medical care in immunocompromised patients, especially in grafted patients. Such opportunistic infections might occur in UC and CD patients as well, meaning that they too would require special medical attention. In this case, the use of vaccines and antiviral therapies should potentially be considered in patients diagnosed with IBD.

## MATERIAL AND METHODS

### Patient and tissue specimens

Tissue samples from individuals with refractory UC and CD who underwent bowel resection between 2010 and 2014 were selected from the Pathology Department files of the University Hospital of Dijon. Approval for the study (reference 18.11.29.52329) was granted by the French national ethics committee (CPP19002), and consent for further histological analysis and *FUT2* genotyping was obtained from each patient. Slides were reviewed by two pathologists in order to rule out dysplasia and cancer according to previously defined criteria (49). Twenty-five surgical resection specimens from 24 IBD patients suffering from either UC (*n*=9) or CD (*n*=15) were selected (Table 1). The patients all presented a clearly defined histopathological and clinical diagnosis. The UC cohort was composed of four males and five females aged 26 to 58 years (median age: 40 years). For the nine patients suffering from UC, samples of distal colon (*n*=6) and rectum (*n*=3) were used for histological analysis. The CD cohort was composed of five males and 10 females aged 22 to 50 years (median age: 32 years). For the 16 CD cases, the samples consisted of jejunoileum (*n*=7, 2 jejunum and 5 ileum), proximal colon (*n*=3) and distal colon (*n*=6, 2 sigmoid colon and 4 transverse colon).

For each case, one block of formalin-fixed paraffin-embedded (FFPE) tissue was selected. For one CD patient, two different surgical resection specimens were retrieved (one block of jejunum and one block of distal colon).

### Histological preparation and antigen detection

Ultra-thin (4 μm) microtome sections were cut from paraffin blocks. Sections were then deparaffinized and rehydrated in xylene and pure ethanol using a Tissue-tek^®^ Prisma^®^ slide stainer (Sakura Finetek Europe). Endogenous peroxidase activity was inhibited using 3% H2O2 in molecular-grade methanol (Sigma-Aldrich, Germany). Slides were washed with phosphate buffered saline (PBS) for 5 min before incubation with 1% bovine serum albumin (BSA) and 3% normal horse serum diluted in PBS, for 2h at room temperature.

HuNoV virus-like particles (VLPs) and antibodies were all diluted in PBS with 1% BSA. Primary monoclonal antibodies (mAbs) were all detected with HRP-labeled mouse specific antibodies (Vector Labs, USA), and incubated for 45 min at room temperature. Peroxydase activity was revealed with 3,3’-Diaminobenzidine for 1.5 min at room temperature (Vector Labs, USA), and sections were rinsed and counterstained with hematoxylin (Dako, Agilent Technologies, USA). From 1 to 5μg of in-house purified GII.4/2007-Osaka (Cairo 4 variant strain (EU876884); hereinafter referred to as “GII.4 VLPs”) and ΔD373 GII.4/2004-Hunter VLPs (E1057 mutant variant unable to bind to HBGAs (EU876890) used as a negative control; hereinafter referred to as “ΔD373 VLPs”) were used for the histological binding assays (37). Production and purification of recombinant VLPs as well as VLP binding assays on histological sections have been described previously (34, 37) (Fig. S6). In-house GII.4-specific mAb directly labeled with peroxydase was used for the detection of GII.4 and ΔD373 VLPs. A and B antigens were detected with 1000-fold diluted mAbs 9113D10 and 9621A8 (Diagast, Loos, France), respectively, whilst H antigen was detected with 1 μg/ml *Ulex Europaeus* Agglutinin I (UEA-I) lectin labeled with fluorescein isothiocyanate (Sigma-Aldrich, Germany). Le^a^ and Le^x^ were detected with 0.5 μg/ml of mAbs MEM-158 and 7Le (Sigma-Aldrich, Germany), respectively. Sialyl-Lewis a and Sialyl-Lewis x (sLe^x^ also known as CD15s) were detected with 2 μg/ml of mAbs NS-1116-19.9 (Dako, Agilent Technologies, USA) and CSLEX1 (Becton Dickinson, USA), respectively. The cell proliferation marker Ki-67 was detected with mAb MIB-1 (Dako, Agilent Technologies, USA). For Ki-67 and sLe^a^ antigens, the epitopes were unmasked by heating at 95°C for 30 min prior to detection on a Dako OMNIS™ automate (Agilent Technologies, USA). When indicated, sections were first treated with 3 mU/ml of *Vibrio Cholerae* sialidase (Sigma-Aldrich, Germany) prior to incubation with antibodies or VLPs, as described previously (34). α1,2 Fucosidase enzyme was a kind gift of Takane Katayama (Kyoto University, Japan) and was used as described previously (34, 50).

For the immunohistological characterization of the tissue, the proportion of labeled area was determined for each sample. The values correspond to the ratio between labeled mucosa and total mucosa of the tissue section as shown in supplemental tables 1 and 2.

### Competition assays

For the characterization of the ligands recognized by the GII.4 HuNoV, histological blocks embedded in paraffin were selected from two patients with blood group A: a 26-year-old woman (sample 13) suffering from CD with a total colectomy, and a 44-year-old woman suffering from UC with sub-total colectomy (sample 22). A tissue area of one square millimeter, comprising regenerative mucosa, was dissected from the original samples for the competition experiments. Sections were first treated with sialidase as described above. Then the sections were incubated with 10 μg of *Helix pomatia* (HPA) and UEA-I lectins (Sigma-Aldrich) in 400 μl/section of PBS overnight at 4°C. After 3 washes with PBS, the sections were incubated with 10 μg of *Aleuria aurantia* or *Lotus tetragonolobus* lectins (all from CliniSciences, France) or both combined, diluted in 400 μl/section of PBS.

For the competition assays using mAbs, sections were either preincubated with 40 μg/ml of Le^x^- or Le^a^-specific mAbs or a combination of both. The sections were then rinsed three times in PBS prior to incubation with 3 μg/ml of purified GII.4 VLP at 4°C for 18 h. VLPs were detected as described above. Treatment of the sections with 50 mM sodium periodate (Sigma-Aldrich) was used for the removal of carbohydrates, as described previously (34).

### FUT2 genotyping

For each patient, DNA was extracted from healthy FFPE tissue. Tissue sections were incubated with proteinase K for 18h at 56°C prior to extraction on the QiaSymphony (Qiagen, USA) following manufacturer recommendations. The DNA was diluted in water and used for the PCR amplification of a portion of the FUT2 gene and sequencing (51). Genetic analysis of the G428A and A385T mutations was performed using the codon aligner suite (CodonCode Corporation, USA).

## ACKNOWLEDGMENTS

This work was supported by the National Reference Center for Viral Gastroenteritis and the Dijon-Bourgogne University Hospital (France). Georges Tarris received a fellowship from the Faculty of Medicine, University of Burgundy (Dijon, France). We would like to thank Suzanne Rankin and Stephanie Lemaire for technical and editorial assistance.

## Data availability statements

Digitized images (WSI format) from the histological analyses and HuNoV VLPs used in the manuscript are available upon request.

## Supplemental material legend

**Table S1:** Summary of HuNoV VLP GII.4 and HBGA detection for CD samples.

For the HES stain section, proportions of mucosal surface areas, quiescent mucosa (QM) or regenerative mucosa (RM) are indicated in parentheses for each patient sample. The information corresponding to RM is shadowed in grey. In the absence of pan-mucosal staining (PM), the specific staining profile is indicated as goblet cell (GC), basal crypt compartment (BC) and epithelial cell (EC).

**Table S2:** Table summary of HuNoV VLP GII.4 and HBGA detection for UC samples.

For the HES stain section, the proportions of mucosal surface areas, quiescent mucosa (QM) or regenerative mucosa (RM) are indicated in parentheses for each patient sample. The information corresponding to RM is shadowed in grey. In the absence of pan-mucosal staining (PM), the specific staining profile is indicated as goblet cell (GC), basal crypt compartment (BC) and epithelial cell (EC).

**Fig. S1:** Ki67 detection in CD and UC samples. Ki67 staining is characterized by dark punctuated staining of the nucleus (arrowhead). Anatomical sites are indicated on the left. Sample origin, Quiescent (QM) and Regenerative Mucosae (RM) are indicated above the panels.

**Fig. S2:** VLP-HBGA interaction in a healthy duodenal biopsy from patient 22 with blood group A. Pretreatments are indicated above the panels. VLPs were directly incubated on non-treated tissue in the mock assay (positive control). Antigen A-specific lectin HPA was used for the competition assays. HPA was inactivated by boiling and used for control. GII.4 VLP specific binding is indicated by arrowheads. Magnified areas are indicated by dashed boxes.

**Fig. S3:** GII.4 VLP inhibition experiments. Sections from UC patient 22 were first incubated with a combination of lectins (LTL, UEA-I and HPA) and sialidase prior to GII.4 VLP binding assays. Sections without treatment or preincubated with boiled LTL were used as controls. Pretreatments are indicated above the panels with plus (treatment) and minus signs (no treatment). VLP binding is indicated by brown staining (arrows). It can be noted that VLP detection showed greater intensity after the removal of sialic acid moiety with sialidase, leading to reduced VLP steric hindrance.

**Fig. S4:** GII.4 VLP inhibition experiments using Aleuria Aurantia Lectin (AAL). VLP inhibition experiments in CD (Patient 13) and UC (Patient 22) slides. GII.4 VLPs were incubated following incubation with Aleuria Aurantia Lectin (AAL). Pretreatment of each tissue section is indicated above. For all experiments, bound GII.4 VLPs (arrowhead) were detected using GII.4 VP1-specific MAb conjugated with peroxydase. Magnified areas are indicated by dashed boxes and shown under each panel.

**Fig. S5:** GII.17 and GII.3 interactions. Attachment of GII.17 and GII.3 VLPs on healthy duodenum (control) and sigmoid colon from one UC resection specimen (patient 22). GII.3 and GII.17 were detected with genotype-specific immune rabbit serum. Goat peroxydase-conjugated serum raised against rabbit IgG were used for the detection of the primary antibody. The VLP genotype is indicated on the left side of each panel.

**Fig. S6:** Baculovirus-expressed purified VLP. One to four micrograms of purified VLP were resolved with MOPS buffer on a NuPAGE gel in denaturing conditions (lanes 2 to 5). Genotype of each VLP is indicated above the gel and Genbank number is indicated in brackets. GII.3 and GII.17 VLPs are mentioned in the discussion and were used in the experiments depicted in Figure S5. Molecular weights in kDa (Lane 1) are indicated on the left side of the gel.

